# Statistical factorization in motor control: Opposing biases from statistics on different timescales enable fast and accurate behavior

**DOI:** 10.64898/2026.07.29.741609

**Authors:** Tianhe Wang, David Whitney

## Abstract

Motor planning is fundamentally constrained by the speed–accuracy tradeoff: a single optimization that balances time and error costs, such that longer preparation yields more accurate movements while faster ones incur greater error. Here we propose that the motor system exploits environmental statistics to reshape this relationship, using two complementary computations that rely on distinct statistical information. The first leverages the marginal statistics of the target distribution, learned by integrating slowly over many trials, to modify the initial preparation state, accelerating planning for frequently encountered targets; this simultaneously speeds decisions and reduces planning variability, but introduces an attractive bias toward the mean of the target distribution. Strikingly, a second process tracks the sequential statistics of the environment over only the last few trials, producing a repulsive sequential bias whose persistence is tuned to the temporal autocorrelation and selectively counteracting the attractive bias introduced by the first. These results reveal that the motor system applies statistical factorization, estimating marginal and sequential statistics on their own timescales to shape movement preparation, thereby escaping the speed–accuracy tradeoff and producing movements that are both faster and more accurate.

## Introduction

Adaptive movement is widely assumed to face a critical tradeoff. While most daily tasks require movements that are both fast and accurate, improving one usually comes at the expense of the other^1–4^. For example, reaching in haste to tap a button on a phone, we hit the wrong one far more often than when we take a moment to aim. In motor planning, this speed–accuracy tradeoff (SAT) is classically attributed to limited neural resources that impose a fixed ceiling on planning efficiency, such that accuracy can be bought only by spending more preparation time^5–7^. With less time to prepare and refine the motor plan, the motor command becomes noisier, inflating endpoint variability and degrading accuracy^8–10^. The same principle extends beyond movement to perceptual decision-making, where committing to a choice sooner integrates less evidence, so speed is again purchased at the cost of accuracy^11–14^.

Classic motor-control theories regard the SAT function as a relatively rigid constraint, one that can be shifted only by recruiting additional neural resources or adopting a more efficient control policy^1,3,10,15,16^. Here we propose a simpler alternative that the motor system can simultaneously raise speed and accuracy by configuring its movement preparation state in advance, according to a prediction of the upcoming movement. Importantly, such a prediction needs to be built from regularities in the world, requiring the motor system to learn and exploit the statistical structure of its environment.

Consider an extreme case in which the same target appears on every trial, a highly predictable environmental statistic. Because the motor system can reliably anticipate this target, we hypothesize that it shifts its initial preparatory state toward the state encoding that movement (Fig 1a)^17,18^. When the target reappears, the system responds both faster and more accurately, moving the SAT function inward within this constrained regime (Fig 1b). Once the environment changes, however, that same anticipatory shift may turn into a cost; with the preparatory state already biased toward the expected movement, the reach toward the new target would be drawn toward the predicted one, producing a systematic attractive error. This is precisely what is observed as use-dependent learning (UDL), a well-documented phenomenon where reaches are biased toward recently repeated movements^19–21^. Interestingly, at the opposite extreme, where every target is drawn at random and is fully unpredictable, movement history exerts the reverse influence. Rather than being pulled toward preceding movements, the current reach is pushed away from the one just made, known as a repulsive sequential effect^22^. As such, opposing biases have been observed across two different environmental statistics, with attraction when the environment is strongly repetitive and repulsion when fully unpredictable.

**Figure 1.**
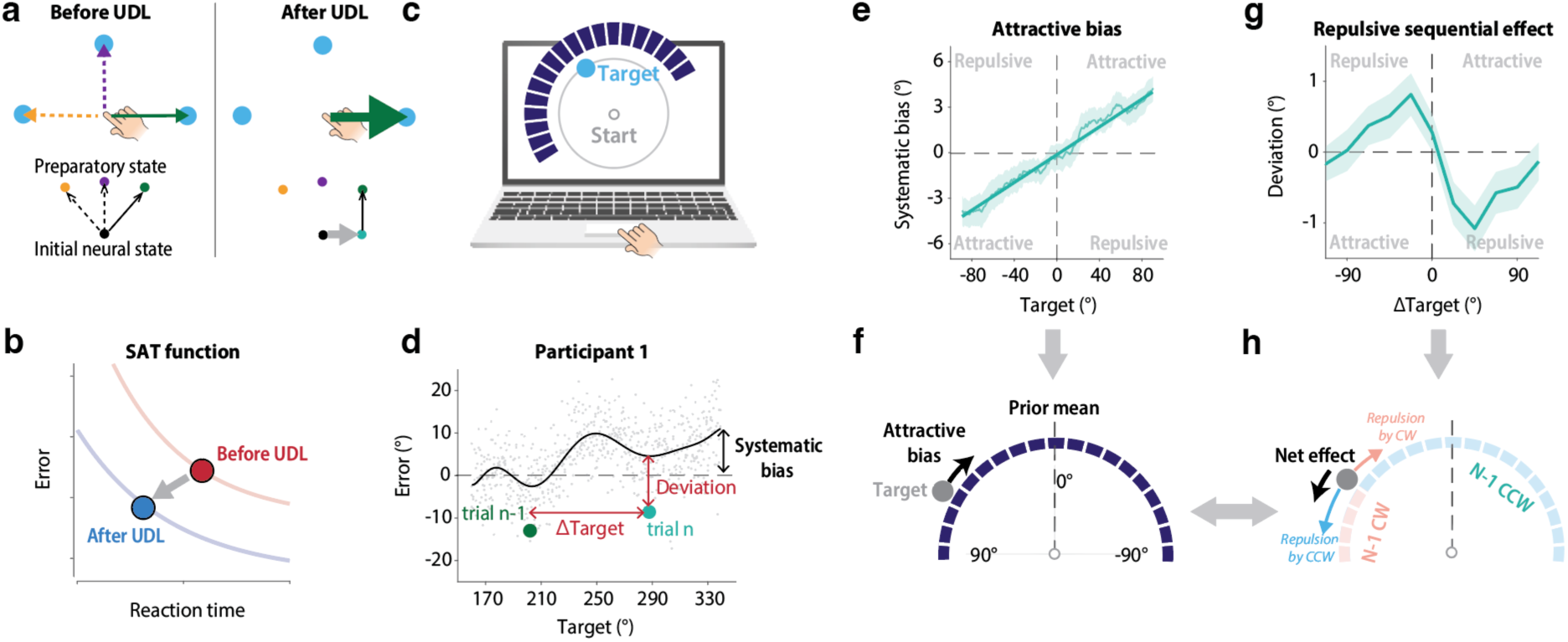
Coexistence of opposite history-driven biases in a center-out reaching task. (a) Schematic illustration of how an expectation of the upcoming sensory input, such as the target location, may facilitate motor preparation in UDL, where participants repeatedly move to a fixed target. Left: during preparation, the neural system starts from a common initial state and transitions to the preparation state that encodes the movement toward the current target. Right: in UDL, the same target is repeated on every trial, so the system can shift its initial state in anticipation of that fixed target, improving preparation efficiency for it. (b) In principle, UDL could reduce preparation time while increasing accuracy when the environment remains unchanged, effectively shifting the SAT function inward. (c) Task setup for Exp 1. Participants made center-out reaching movements using a trackpad while targets and a cursor were displayed on a laptop monitor. Blue bars indicate the semicircular uniform distribution from which reach targets were sampled. (d) Raw reaching error from a representative participant. Gray dots indicate individual trials. The black curve shows a polynomial fit capturing systematic bias, reflecting a combination of target-specific bias and central tendency. Deviation is defined as the trial-by-trial difference between the observed reach and this systematic bias function. (e) Attractive bias toward the mean. The thin-blue curve shows the group-averaged systematic bias, revealing attraction toward the center of the distribution. The thick-blue line indicates a linear fit. Shaded regions indicate ±1 s.e.m. (same below). (f) Illustration of the attractive bias for a given target. (g) Repulsive sequential effect. Deviation is plotted as a function of ΔTarget (defined in d). Reaches are biased away from the previous target when ΔTarget is within ±90°. (h) Illustration of how trial-by-trial repulsive effects generate a bias that offsets central tendency when averaged across trials. For a target located at 50°, the previous target is more likely to have appeared in the counterclockwise (CCW) direction relative to the current target than in the clockwise (CW) direction. Consequently, repulsive sequential effects occur more frequently from the CCW side, producing a net repulsive force that compensates for the attractive central tendency.

How does the motor system balance these opposing biases? What computation produces them, and how does it exploit environmental statistics? While previous work explored motor biases under extreme statistics, natural environments are much richer and usually fall somewhere in between the extremes. A tennis player varies the placement of every serve, and a typist moves continually across the keyboard; there is some repetition, but also some unpredictability or randomness. In such situations, the opposing biases could arise from a single mechanism that tracks the volatility of the environment, grading continuously from attraction in stable surroundings to repulsion in variable ones. Alternatively, the opposing biases might reflect two dissociable computations that factorize the statistics of the environment and run in parallel on different timescales: a slow one that reads the marginal statistics (the overall target distribution) to set the preparation state, and a fast one that reads the sequential statistics (the dependence between successive targets) and continuously cancels the attractive bias the slow process introduces. Such a division of labor could keep the speed gained from anticipating the overall structure while offsetting the error it introduces, rather than trading one against the other.

To distinguish these possibilities, we employed a center-out reaching task while manipulating target distributions and target autocorrelation, creating environments with different predictability and temporal dynamics. We examined their joint effect on planning speed, overall accuracy, and bias patterns. Coupled with computational modeling of motor planning, our results suggest that the attractive and repulsive biases operate simultaneously based on environmental statistics on different timescales. Together, our results reveal statistical factorization in motor control, an architecture that enables movement to be both fast and accurate.

## Results

### Coexistence of attractive and repulsive influences from movement history

To understand how motor planning exploits environmental statistics, Exp 1 was designed to identify the bias structure in a motor control task with variable target positions. Participants made center-out reaches toward targets randomly sampled from a uniform distribution covering a semicircle (Fig 1c, S1). We measured angular error in the movement as participants crossed the target circle. Consistent with previous reports, we found a systematic attractive effect toward the prior mean (Fig 1e), in the direction predicted by UDL^20,21^. Critically, by analyzing trial-by-trial fluctuations of the reaching angle (the deviation index, Fig 1d) as a function of the difference between current and previous targets (ΔTarget), we observed a repulsive sequential effect: the current movement was biased away from the immediately preceding one (Fig 1g)^22^. While both effects have been reported independently in previous work, those studies employed either a highly constrained or a completely random target distribution, allowing only one of the effects to be tested in isolation^20,22–24^. The current results demonstrate that both effects occur simultaneously under a naturalistic target distribution, suggesting they reflect separate mechanisms adapting to statistics on different timescales.

This pattern of opposing biases raises a fundamental question: what mechanism drives these biases? Environmental statistics vary along two dimensions. The marginal statistics describe the distribution from which targets are drawn, irrespective of their order; the sequential statistics describe the dependence between successive targets. We hypothesize that the two effects reflect distinct computational processes, each reading one of these statistics. One generates an expectation of target position from the marginal statistics, speeding movement preparation at the cost of an attractive bias; the other draws on the sequential statistics to provide a distributed correction that counteracts this attractive bias. We next develop theoretical analyses of each proposal.

### An adapted drift–diffusion model for movement preparation

We propose that the attractive bias observed in Exp 1 arises from a process in which the motor system adjusts its preparation state based on task-specific priors. To illustrate this mechanism, we developed a computational framework based on a spatial drift–diffusion model (sDDM)^11,25^ that adapts to the target prior distribution (Fig 2a, left). The sDDM comprises a neural population tuned to the full 360° range of potential movement directions^26,27^. Upon target presentation, evidence for the corresponding movement accumulates over time as a Gaussian-shaped function centered on the target, with each unit bearing independent noise^28–30^. A movement is initiated once the accumulated evidence of the peak unit surpasses the decision boundary. This model generates a typical SAT function when the decision boundary is varied: raising the boundary allows more time for evidence accumulation, boosting the signal-to-noise ratio and increasing planning accuracy (Fig 2a, right).

**Figure 2.**
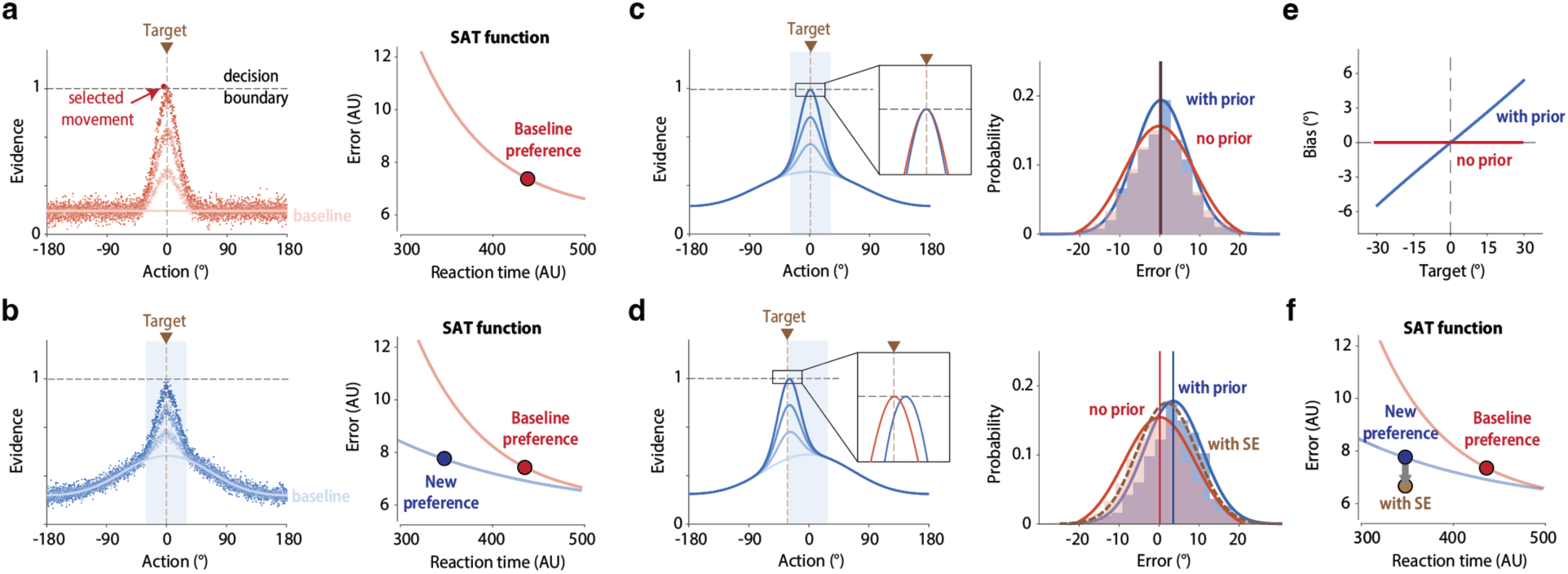
A spatial DDM illustrating how learning a target distribution shifts the speed–accuracy tradeoff function. (a) Schematic of a spatial DDM variant implemented with discretized population units representing movement directions. *Left*: Upon target presentation, evidence accumulates around the target direction. The action corresponding to the first unit to reach the decision boundary is executed. Due to drift noise, the selected action may deviate from the target. *Right*: The speed–accuracy function, generated by systematically varying the decision boundary to evaluate root-mean-square error (RMSE) at a given reaction time (RT). The dot indicates the operating point corresponding to the decision boundary shown on the left. (b) Same as (a), but for a condition where targets are sampled from a restricted 60° range (shaded area). The system modulates its initial state to reflect this prior, shifting the SAT function inward. (c) Model response to a target at the prior mean. *Left*: Expected evidence accumulation for the center target after learning the 60° prior. The cross-section shows the state profile at the moment the peak unit reaches the decision boundary, comparing the full model with the learned prior (blue) to a lesioned model assuming no prior (red). The two models are compared at a matched RT. The lesioned model exhibits a slightly broader state profile. *Right*: Consequently, the lesioned model produces greater response variance in the presence of drift noise. (d) Model response to a target at −30° (the edge of the prior). While the full model maintains narrower variance, it systematically shifts the response mean toward the prior mean because the initial state profile reflects the shape of the prior distribution. Moreover, the repulsive sequential effect pushes in the opposite direction to this attractive bias and further increases accuracy (yellow dashed line). (e) Prior learning introduces an attractive bias toward the mean which increases in a close-to-linear manner. (f) The repulsive sequential effect can counteract the attractive bias, further reducing error without incurring an RT cost (labeled “with SE”).

Importantly, we assume that the system adapts to the target prior by shifting the movement preparation state (Fig 2b, left), such that units tuned to directions near the prior mean start from a higher initial state, and are therefore closer to the decision boundary. This modulation has three consequences. First, less additional evidence is required for targets near the prior mean, shortening preparation time. Second, this inhomogeneous initial state also reduces the decision variability for central targets by compressing responses toward the prior mean (Fig 2c). Third, as a byproduct, the response shows a systematic bias toward the prior mean (Fig 2d), one that grows as the target diverges further from the center (Fig 2e). Taken together, the adapted preparation state increases preparation speed and reduces variability, effectively shifting the simulated SAT function inward for the learned environment despite the accompanying central-tendency bias (Fig 2b, right).

Moreover, we propose that the repulsive sequential effect can serve as a compensatory mechanism that reduces this central-tendency bias and further increases planning accuracy. Specifically, as shown in Fig 1g, the trial-by-trial repulsive bias provides a distributed correction signal. For a given target offset from the prior mean, the preceding target that drives the repulsive effect is more likely to fall on the central side than on the edge side. Consequently, when marginalized across all possible previous target locations, the repulsive effect produces a net bias opposite to the central tendency (Fig 2f). Under this view, the motor planning system modulates the SAT function through the cooperation of two separate processes, each exploiting distinct environmental statistics on a different timescale. We tested these proposals in the experiments that follow.

### Exploiting the target distribution increases planning efficiency

To test the proposed mechanisms, we start by focusing on how learning the target distribution modulates the SAT function. In Exp 2 we examined the co-modulation of motor bias and RT using the same center-out reaching task as Exp 1, but this time targets were sampled from one of three distributions (Fig 3a): a 180°-range (wide, i.e., Exp 1), a 120°-range (medium), or a 60°-range (narrow) uniform distribution. Importantly, to probe the SAT function while allowing participants to express their natural preferences, we used two conditions with different response contingencies. In the delayed-response condition, a separate go cue was presented 700 ms after target onset (Fig S1), so that participants had sufficient time to fully prepare the movement during the delay. In the free-response condition, target onset itself served as the go cue, and participants could initiate the movement as soon as they detected the target (Fig S1). Whereas participants may maintain a stable decision boundary in the delayed-response condition, they are likely to collapse the boundary over time in the free-response condition, initiating movement sooner at the cost of higher error. These two conditions thus probe distinct points on the SAT function.

**Figure 3.**
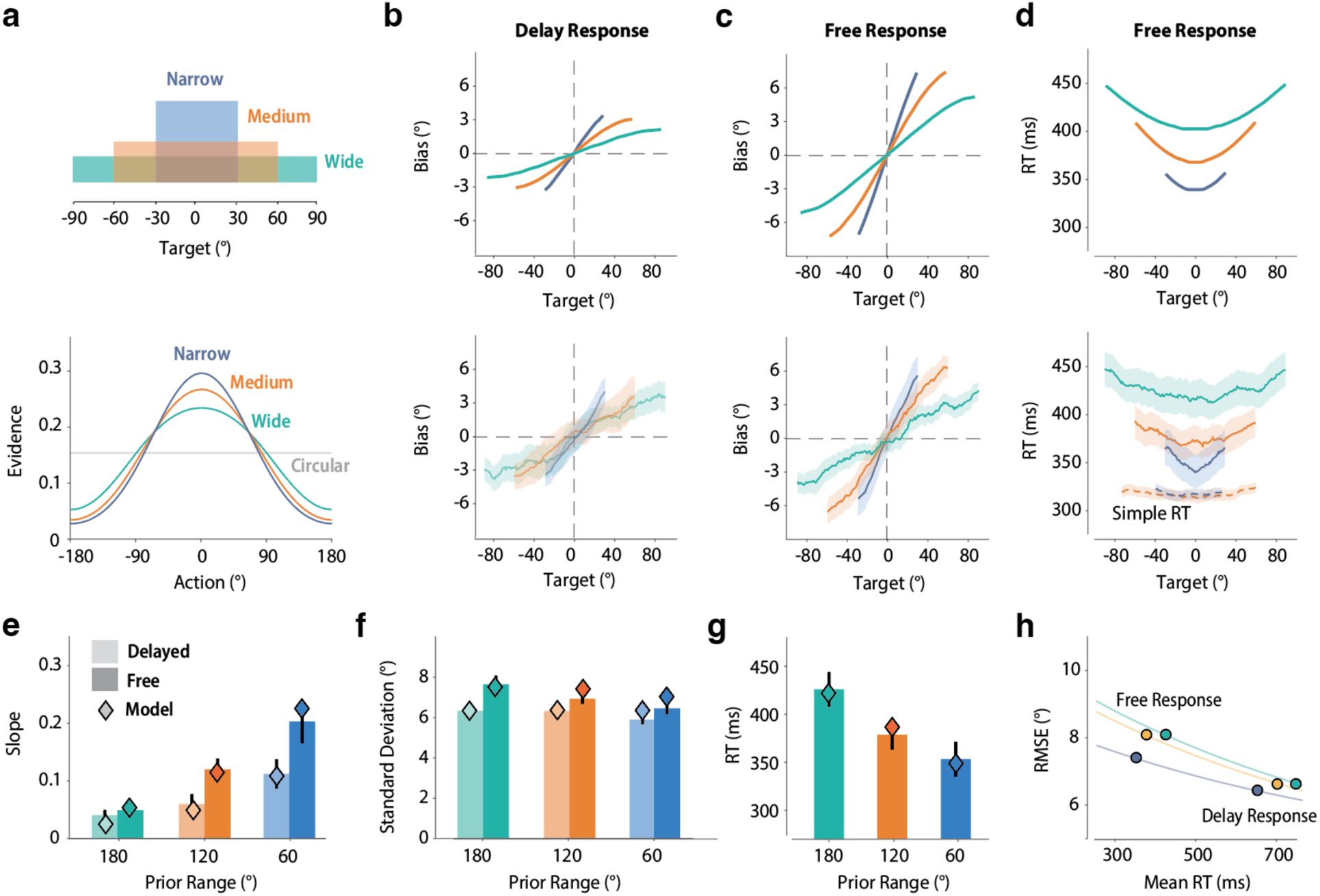
Target prior distribution co-modulates motor bias, variance, and RT. (a) Three target prior distributions of different ranges applied in Exp 2 (top) and the corresponding initial state in the sDDM after learning each prior (bottom). (b–c) The attractive bias function predicted by the sDDM (top) and measured from the behavioral results (bottom) for the delayed-response (b) and free-response (c) conditions. The slope increases as the prior range narrows, and the bias is stronger in the free-response than the delayed-response condition. (d) U-shaped RT function predicted by the sDDM (top) and measured from the behavioral results (bottom) for the free-response condition. RT decreases as the prior range narrows and as the target falls closer to the prior mean. Dashed lines in the bottom panel show the simple-RT results from the control condition (Exp S1, medium and narrow conditions only). (e–g) The slope (e) and standard deviation (f) of the motor bias functions across all conditions, and the average RT in the free-response condition (g), each aligned with the prediction of the best-fitted model (diamonds). (h) RMSE as a function of RT across prior conditions. Free-response allows faster responses at the cost of higher error; a narrower target prior lets the system prepare faster and more accurate movements, shifting the SAT function inward. RT for the delayed-response condition was derived from the sDDM and is plotted here for illustration.

Based on the sDDM, a narrower prior creates a stronger initial-state modulation (Fig 2e), which should generate shorter RTs, lower planning variability, and a stronger central-tendency bias. The model further predicts that both the attractive bias and the motor variability should be larger in the free-response than in the delayed-response condition, because only in the free-response condition does the decision boundary collapse over time, forcing earlier and less-prepared movements.

All of these predictions were confirmed by the behavioral results. Beginning with the attractive bias, both response types produced a close-to-linear central-tendency function whose slope increased markedly as the prior narrowed (F(2,190) = 14.1, p < 0.001, 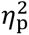 = 0.13, Fig 3b–c). However, the average magnitude of the bias remained comparable across prior conditions, because a narrower prior also confines targets to a smaller range of eccentricities; the error cost imposed by the attractive bias therefore stays limited under a narrower prior. As predicted, the bias was also larger in the free-response than the delayed-response condition (F(1,190) = 8.8, p = 0.004, 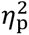 = 0.044), confirming that participants traded accuracy for speed when free to initiate movement.

Turning to the benefits of prior adaptation, motor variability decreased as the prior narrowed (F(2,190) = 3.11, p = 0.047, 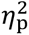 = 0.032). Importantly, we also observed clear RT modulation in the free-response condition (F(2,90) = 4.98, p = 0.009, 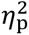 = 0.10). Overall RT fell as the prior range narrowed (Fig 3g), and within each condition RT followed a U-shaped profile, with movements toward targets near the prior mean initiated faster than those toward the edges of the distribution (Fig 3d, ts > 5.1, ps < 0.001). A simple-RT control under the same variable target priors confirmed that this modulation reflects faster movement preparation rather than faster visual detection (Exp S1). In the delayed-response condition, by contrast, RT showed no such modulation (Fig S2a; F(2,100) = 0.30, p = 0.74, 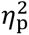 = 0.006), presumably because preparation finished within the 700 ms delay. Movement time, however, showed a U-shaped profile in both response-contingency conditions, fastest toward the prior mean (Fig S2b; ts > 3.2, ps < 0.002), an additional benefit of prior adaptation. Together, these results suggest that prior statistics reduce preparation time without inflating error, thereby shifting the SAT function inward.

We then examined whether the sDDM could quantitatively account for the bias, variability, and RT patterns in Exp 2. Using the delayed-response data, we fitted the bias functions across the three prior conditions to determine the learning rate of the initial preparation state function, the widths of the two generalization functions, the decision bound, the saturation cap on the initial-state field, and the noise (see Methods). Holding these parameters fixed, we then fitted a single boundary-collapse rate to the free-response bias functions. With this parameter set, the sDDM quantitatively reproduced all six bias functions and the motor variability across the two response conditions. Notably, the model also reproduced the RT functions observed in the free-response condition, indicating that the sDDM captures a shared computational mechanism coupling preparation speed and motor accuracy. Finally, we plotted the SAT function across the three prior conditions (Fig 3h), estimating the preparation time for the delayed-response condition from the sDDM. Together, these results demonstrate that the motor system tunes its preparation state to the upcoming target, enabling movement that is both faster and more accurate; this benefit grows as the target distribution narrows and thus becomes more predictable.

We considered several alternative models to explain the bias pattern in Exp 2. Whereas we interpret the attractive bias as a cost of faster responding within the SAT framework, previous motor-control theories instead attribute it to Bayesian integration, which combines a noisy input with a learned prior to produce a movement plan that minimizes overall uncertainty^21,31^. However, the bias pattern in Exp 2 is inconsistent with a Bayesian model (Fig S3), an unsurprising result as the current design carried minimal perceptual uncertainty. Moreover, we examined an alternative sDDM variant in which the prior modulates the evidence-accumulation rate rather than the initial state; it too failed to provide a comprehensive account of the behavioral results (Fig S4).

### Bimodal biases in use-dependent learning arise from a single mechanism

To further test our sDDM, we ask whether the model accounts for phenomena under other statistics, such as UDL, in which a single target is repeated, creating an extremely stable environment. In a typical UDL design, participants repeat a specific movement for hundreds of trials, while the target switches to a novel one unpredictably in probe trials. Interestingly, the error distribution for probe trials shows a bimodal pattern (Fig 4a–b Left). Most reaches land near the probe target with a small attractive bias, while a smaller subset lands near the repeated target itself, forming a second cluster with large errors. Notably, these large-error habitual responses have shorter RTs than the accurate, target-driven responses, so that error and RT are coupled within probe trials. Previous theories interpreted this bimodal pattern as evidence for two distinct processes: an action-selection system responsible for the large habitual errors and an action-execution system responsible for the small biases around the target^23,24^.

**Figure 4.**
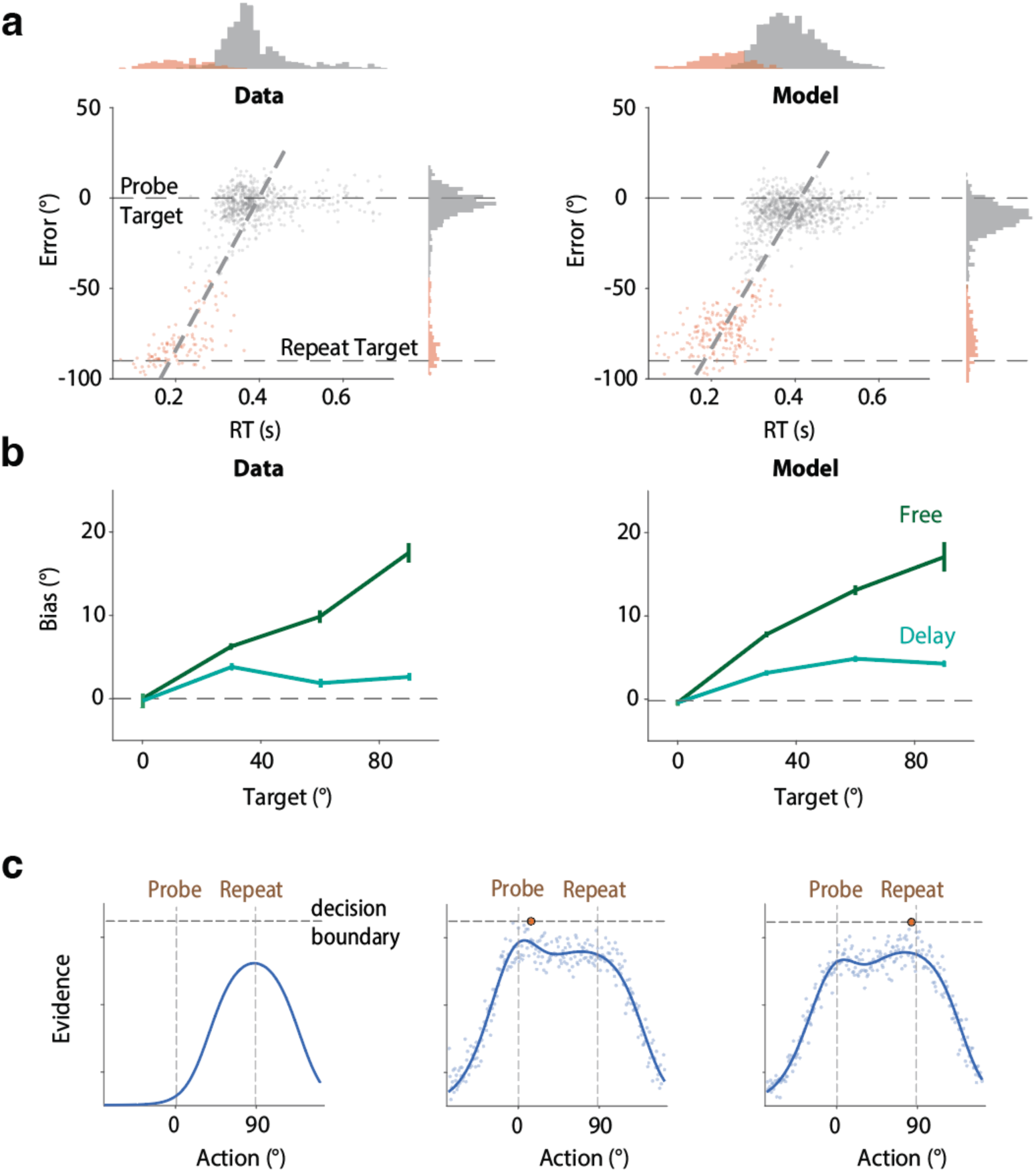
The adapted drift–diffusion model accounts for bimodal error with a single mechanism. (a) Relationship between RT and reaching error in use-dependent learning. Data are replotted from a previous experiment^23^ in which participants repeatedly reached to a center target, with occasional probe trials to a target 90° away. Reaching error and RT both exhibit bimodal distributions (shown in the marginals). For illustration, trials are split by error magnitude and plotted in different colors. The model reproduces both the bimodal distributions and the correlation between error and RT. (b) Mean reaching bias as a function of probe-target distance relative to the repeated target, for both the free-response and delayed-response tasks. (c) Illustration of how the sDDM generates the bimodal bias. Left: the repeated target drives a strong modulation of the initial-state function in anticipation of the repeated movement. Middle: when a probe target appears, evidence accumulates around the probe target, but the adapted initial state introduces a central-tendency bias. Right: on some trials, units near the repeated target reach the decision boundary before evidence accumulation is complete, producing a fast, habitual response with a large error. The dark blue curve represents the noise-free evidence accumulation function; blue dots indicate individual units; the orange dot indicates the boundary-crossing unit.

The sDDM provides an alternative account, predicting this bimodal error without positing dual systems (Fig 4c). Specifically, the system learns a highly concentrated prior after a single target is repeated hundreds of times. When a probe target is presented, stochastic noise can occasionally drive units tuned to the prior peak across the decision boundary before units tuned to the current target, giving rise to a peak with habit-like responses (Fig 4a right). As the two populations compete within a single accumulation process, both modes are skewed inward (Fig 4c). Moreover, as units tuned to the repeated target start from a much higher initial state, they reach the boundary earlier, so habit-like responses have shorter RTs than target-driven movements. Our model thus demonstrates that the bimodal error pattern can emerge from a single drift–diffusion process with learned prior adaptation (Fig 4a–b Right), without requiring separate habitual and goal-directed control systems.

### The repulsive sequential effect reduces the attractive bias

Having shown how prior adaptation speeds up movement preparation while introducing an attractive bias, we next asked whether the repulsive sequential effect serves as a compensatory mechanism that reduces this bias and preserves planning accuracy. To examine this, we began with a model simulation based on Exp 1. We generated reaching angles from a model that expressed either the attractive bias with the repulsive sequential bias, or the attractive bias alone with no sequential bias (Fig 5a). RMSE was smaller in the former case than in the latter, suggesting that the repulsive effect improved reaching accuracy in Exp 1 (Fig 5c). Expanding the simulation to a range of sequential-effect strengths, we found that error followed a U-shaped function of this strength (Fig 5e): increasing the sequential effect reduces the attractive bias but also raises trial-to-trial response variability, so overall error is minimized at an intermediate strength (Fig 5d). Notably, the sequential effect observed in Exp 1 falls close to the valley of the function where the maximal accuracy is achieved (Fig 5e).

**Figure 5.**
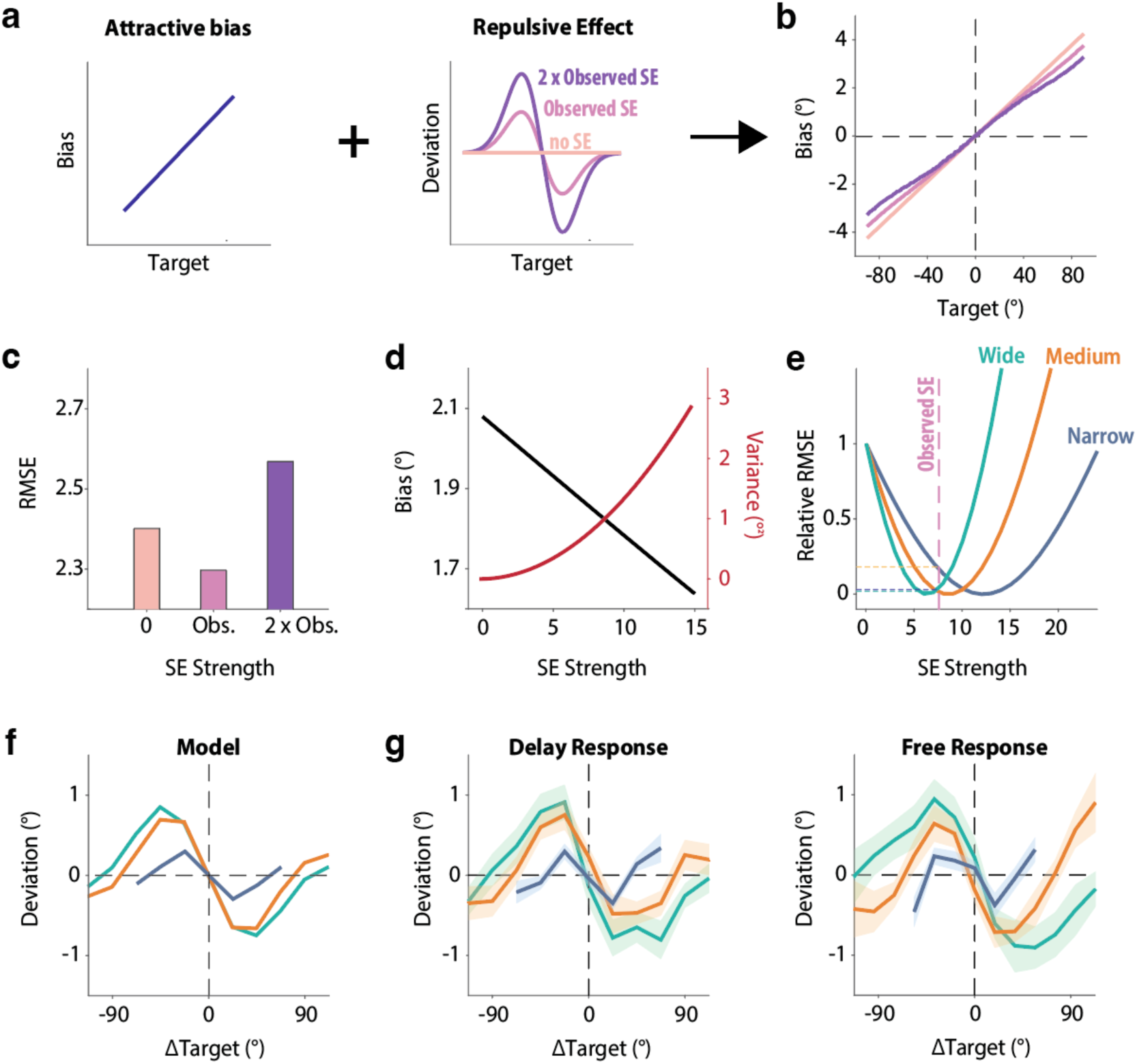
Repulsive sequential effects compensate for central tendency bias. (a) Simulation combining central tendency in the wide prior condition with sequential effects of varying strengths. (b) Increasing the strength of the sequential effect leads to a corresponding decrease in central tendency. (c) Simulations using the observed sequential effect generate smaller RMSE compared to simulations with either no sequential effect or a doubled (2×) sequential effect. (d) Changes in bias and variance as a function of sequential effect strength for the wide prior condition. (e) Relative RMSE as a function of sequential effect strength for wide, medium, and narrow prior conditions given the central tendency measure in the delayed-response condition in Exp 2. The observed sequential effect is recovered from the wide prior condition (see panel f); this effect compensates for the majority of relative RMSE across all three prior conditions. (f) Recovered sequential effects simulated from a model assuming a constant sequential effect across all priors. The sequential effect is strongly distorted as the prior becomes narrower. (g) Sequential effect functions observed across different prior conditions and response contingencies.

A natural question is whether the repulsive effect scales with the prior range to match the growing attractive bias under narrower distributions (Fig 5e). Empirically, however, this is difficult to test directly, because for a narrow prior ΔTarget becomes highly correlated with absolute target position (Fig S5a), confounding the measurement of the sequential effect with central tendency (Fig 5f). To circumvent this, we simulated the expected measurement of the sequential effect across prior ranges under the assumption that the latent repulsive effect remains constant. These simulated curves matched the observed data well, suggesting that the repulsive effect may not scale with prior range. It was likewise unaffected by response contingency (main effect: F(1,190) = 1.5, p = 0.23, 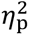 = 0.008; interaction: F(2,190) = 1.4, p = 0.24, 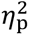 = 0.015), showing a comparable magnitude in the free-response and delayed-response conditions (Fig 5g). This is anticipated as the repulsive effect is mostly useful for offsetting the attractive central-tendency bias rather than the large errors of fast habit-like responses. We then evaluated the compensatory benefit of the repulsive effect across prior conditions for delayed response. Note that even a fixed repulsive effect compensates for more than 80% of the achievable RMSE reduction across all prior ranges (Fig 5e). As such, it is likely that the repulsive effect is tuned to maximize accuracy across a broad range of target distributions but, as a trial-by-trial process operating on a short timescale, is insensitive to the marginal statistics of any specific environment (i.e., the prior range) that required a long-term integration.

### Repulsive sequential bias is enhanced in autocorrelated environments

The results above establish a dissociation that the repulsive effect, presumably relying on sequential statistics, is blind to the marginal statistics that drive the attractive bias. We next asked whether it is instead governed by the temporal autocorrelation of recent targets, the standard measure of sequential structure. So far, our experiments used targets randomized across trials. This is ideal for isolating the intrinsic sequential bias, but motor goals in everyday tasks are often autocorrelated in time. As in handwriting or repeatedly reaching for tools on a bench, successive movements tend to evolve smoothly within a local region. In such an environment, a run of nearby targets drags the preparation state toward that local cluster, so the previous target becomes more informative about the running mean and a more useful signal for compensation. UDL is an extreme but useful illustration. A single target repeats on nearly every trial (Fig 6a), so autocorrelation is high and prior adaptation drives attractive bias toward the repeated location. When the environment suddenly changes on a probe trial, the previous target reflects the accumulated prior that the attraction is built on, so pushing away from it is especially useful. Indeed, we observed a repulsive sequential effect when analyzing how a probe trial biases the subsequent repeated trial (Fig 6b). Assuming the same effect operates in the reverse direction, this repulsion would compensate for the attractive bias on the probe trial more efficiently than in Exp 1–2 (Fig 6c; t(48) = 4.1, p < 0.001, 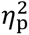 = 0.262).

**Figure 6.**
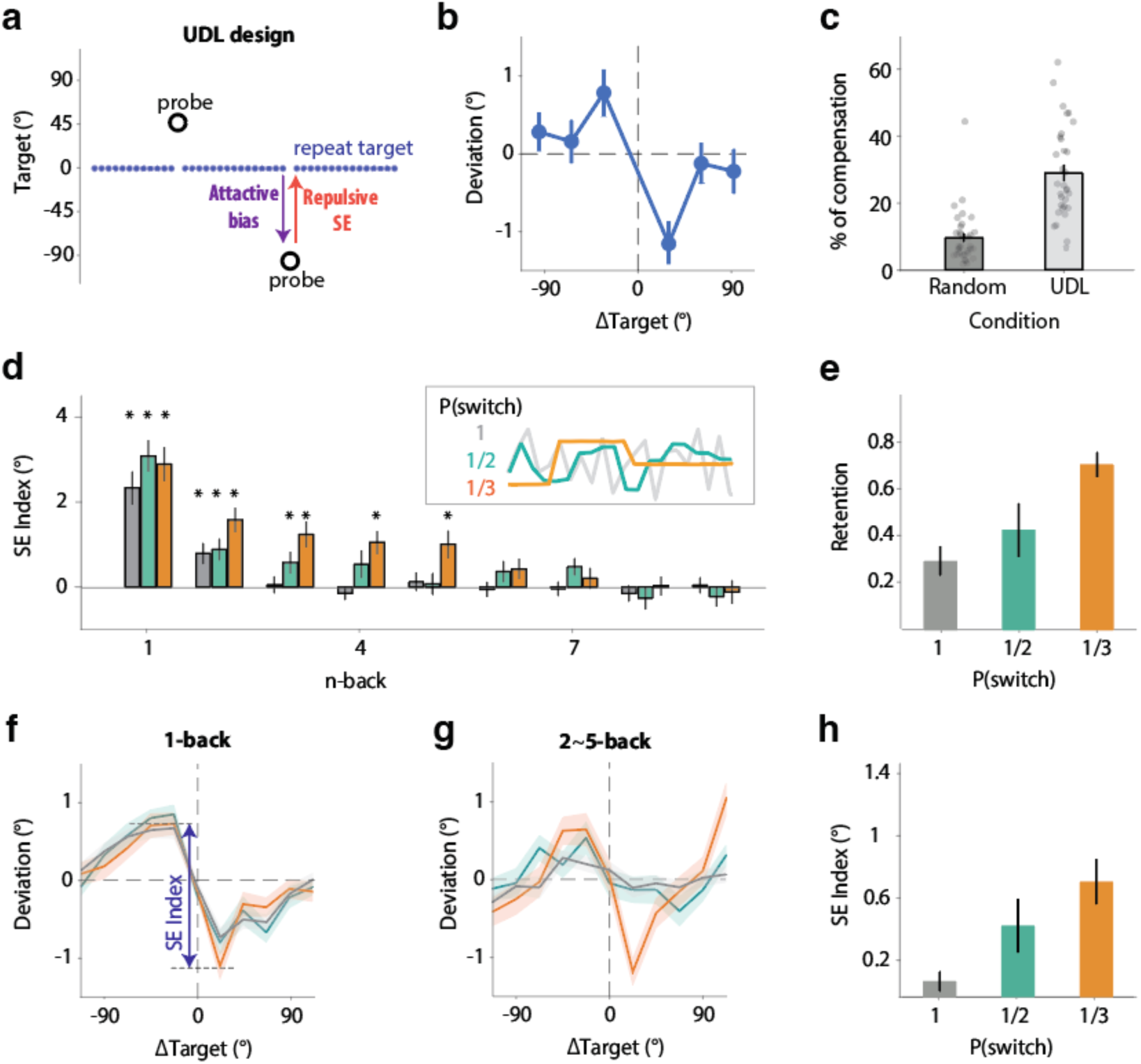
The temporal persistence of the sequential effect is enhanced by environmental autocorrelation. (a) Example target sequence in a UDL design. Participants reached toward the repeated target on 90% of trials, with 10% directed to a probe target. UDL attractive bias is analyzed as the bias at the probe target, while the repulsive sequential effect is measured as how a probe target influences the subsequent reach toward the repeated target. (b) The repulsive sequential effect function observed in the UDL design from a previous study^23^. (c) The percentage of the attractive bias at the non-center target that is offset by the observed sequential effect is higher in the UDL design than with random targets. (d) Strength of the sequential effect as a function of the temporal distance (n-back) between the previous and current trial. The inset shows the target sequence of a 20-trial period for three conditions in Exp 3. The sequential effect persists longer in environments with higher autocorrelation. *: one-sample t-test, p < 0.05 (e) The retention rate (1 − decay rate) of the sequential effect increases as environmental autocorrelation increases. (f) The sequential effect from the 1-back trial remains consistent across conditions with different autocorrelation levels. SE index is the peak-to-peak amplitude of the S-shaped function (see Methods). (g–h) The sequential effect function (g) and SE index (h) averaged across 2-back to 5-back trials. The averaged effect increases significantly in environments with higher autocorrelation.

To further test whether the sequential effect is modulated by temporal autocorrelation, we manipulated the target switching probability in Exp 3 (Fig 6d). Specifically, we compared conditions in which the target switched position on every trial (P(switch) = 1), on 1/2 of trials, or on 1/3 of trials. After each switch, the new target was selected randomly on the circle, so no global geometric mean existed to confound the measurement of the repulsive sequential effect. We restricted our analysis to how the last trial before a switch influenced subsequent movements, as model simulations show that this approach recovers the true sequential effect while avoiding artifacts from autocorrelated movement sequences (Fig S5).

When plotting the sequential effect from the immediately preceding (one-back) trial, we observed similar magnitudes of repulsive bias across all three autocorrelation conditions (Fig 6f; Welch F(2,84.0) = 2.20, p = 0.12). Extending this analysis to include recent trial history, however, revealed distinct temporal dynamics: the sequential effect persisted longer as environmental autocorrelation increased. In the random (P = 1) condition it lasted only two trials, whereas it extended to the third-back trial in the P = 1/2 condition and the fifth-back trial in the P = 1/3 condition (Fig 6d). Fitting an exponential decay to the temporal profile of the effect yielded significantly different decay rates across the three conditions (Welch F(2,69) = 5.9, p = 0.004, Fig 6e) and averaging the effect across the second- to fifth-back trials revealed a clear monotonic increase with autocorrelation (Welch F(2,70.8) = 9.01, p < 0.001, Fig 6g–h). These results demonstrate that the repulsive sequential bias is flexibly tuned by the autocorrelation of the environment, its temporal reach growing as recent history becomes more informative. This makes the correction efficient: the fixed magnitude sits near the value that best offsets the bias without adding excess variability (Fig 5e), while the reach extends over just those trials on which recent movements remain predictive.

Together, these experiments reveal how the motor system partially escapes the speed–accuracy tradeoff: one process adapts to the marginal statistics, increasing response speed at the cost of an attractive bias, while a second, faster process adapts to the sequential statistics and reduces that bias, allowing motor planning to be both fast and accurate across a wide range of environments.

## Discussion

In natural environments, movements must often be both accurate and rapid^32,33^. Whether escaping a predator, catching a falling object, or avoiding an obstacle while walking, success depends on acting quickly while still reaching the correct location^34–36^, creating a fundamental tension in motor planning. Because the neural resources for movement preparation are limited, accuracy is classically thought to improve only by allowing more time to prepare, giving rise to the speed–accuracy tradeoff^1,8,10,13^. Here we show that the motor system can partially escape this constraint by shaping its preparatory state according to the statistics of the environment, through two mechanisms that introduce opposing biases. Prior adaptation reshapes the initial preparation state toward the target distribution, speeding planning and reducing variability but, as a byproduct, biasing movements toward the distribution mean. A repulsive sequential bias, driven by the sequential statistics of recent experience, counteracts this attractive bias and restores accuracy. Because the benefit and its cost arise on different timescales, a slowly learned distribution versus fast trial-to-trial history, the sequential correction removes the bias while preserving the speed and variability gains that prior adaptation produced. Together these mechanisms form a statistically factorized architecture that keeps movement both fast and accurate across a broad range of environments.

### Adapting the preparation state to environmental statistics

Living in a noisy world, the brain learns the statistical structure of the environment to anticipate what is likely to happen next and prepare behavior accordingly^37,38^. This capacity is well documented in perception, where estimates of visual orientation are biased by the statistical regularities of natural scenes^39–41^, and in sensorimotor learning, where participants acquire an imposed distribution of target displacements and rely on it more heavily as visual feedback becomes unreliable^31,42^. The most influential formalization of this ability is Bayesian inference^28,31^, in which a learned prior is combined with noisy sensory evidence to minimize uncertainty. The same idea has been applied to motor control to explain UDL^21,43^, the tendency for movements to drift toward frequently repeated actions^19,20^, where repetition sharpens a prior over movement direction that, combined with the current target, produces an attractive bias toward the frequent direction.

However, several lines of evidence argue against the Bayesian account in motor planning. The graded, near-linear bias that originally supported it^21^ was later shown to be an artifact of averaging over a bimodal distribution, in which most probe reaches land near the true target and a minority near the repeated location, so their mixture mimics a smooth central tendency^23,24^. Once the motor-related component is isolated, the bias pattern departs from the Bayesian prediction (Fig S3). This failure is unsurprising theoretically. Bayesian integration recovers a hidden state from noisy input and helps when the movement outcome is uncertain, as when hand feedback is displaced^28,29,31^, but in an unperturbed natural environment the target carries almost no state uncertainty, leaving little to infer. The challenge here is instead to prepare the movement efficiently under time pressure, a distinct computational goal that calls for a different use of environmental statistics.

Our model offers a more parsimonious account of UDL. The bimodal errors, previously attributed to two separate systems^23,24^, instead arise from a single computation. Repetition raises the initial preparation state of units tuned to the habitual direction, which within a drift–diffusion process yields both the graded attractive bias and the fast, short-latency habitual errors. In this view, UDL reflects an exceptionally stable statistic that drives a strong prediction and heavily modifies the initial state. The same mechanism extends to the more graded statistics of natural environments, where we showed that it reduces both reaction time and motor variability across a range of prior distributions.

What is the potential neural basis of prior adaptation in motor planning? Although our model is largely descriptive, its architecture, a population of units tuned to different movement directions competing to cross a decision threshold, resembles proposed cortico-basal-ganglia circuits for action selection, in which the basal ganglia implement an optimal, accumulation-to-threshold decision among alternative movements^44–46^. Whether the initial state of this circuit is modulated as our model posits remains open for future test. Movement preparation has been characterized as cortical population dynamics that set the initial state for the upcoming movement^47–49^. Consistent with the idea that this preparatory state is tuned to environmental statistics, a recent study manipulated the probability of the direction of an upcoming mechanical perturbation and found that the preparatory population state in dorsal premotor and primary motor cortex shifted systematically, its position in neural activity space scaling with how likely each direction was^17,18^. Our model extends this principle from a discrete, binary case to the graded statistics of a continuous space and characterizes its concrete behavioral benefit of faster and less variable movement preparation. How is this preparatory bias acquired? Recent work has identified a candidate teaching signal. The tail of the striatum carries a value-free, movement-based dopaminergic action-prediction-error that reinforces recently performed actions independently of their value^50^, well suited to gradually sculpt the prior over movement direction assumed by our model.

### Factorizing marginal and sequential statistics across timescales

In a natural environment, the marginal statistics^42,51,52^ and sequential statistics^53–55^ of movement goals can vary relatively independently. Repeatedly reaching for the same tool on a bench couples a narrow range with high autocorrelation, whereas returning a series of varied tennis serves couples a wide range with low autocorrelation. Our results show that two mechanisms adapt to these statistics separately and are modulated by different variables, with the attractive bias scaling with the range of the marginal distribution and with response contingency (i.e., preparation time), and the repulsive bias tuned to the temporal autocorrelation of recent targets. Each dependency is well matched to the statistic its system exploits. The marginal distribution is a stationary, aggregate property, so estimating it well requires integrating over a long stretch of experience until trial-to-trial fluctuations average out, yielding a stable prior that tracks the shape of the distribution and is insensitive to the order of recent targets^51,52^. The sequential dependence is instead a local property, informative only about the immediate past and exploitable only by weighting the most recent trials heavily and forgetting them quickly, so the sequential effect tracks how far into the past recent movements remain informative^53–55^. The system therefore benefits from statistical factorization, pairing a slow integrator that accumulates the marginal distribution with a fast, recency-weighted process that tracks the sequential dependence. Because the two systems rely on different information about the environment, the fast correction cancels the attractive bias without disturbing the slowly learned prior, increasing accuracy without compromising the speed benefit.

Interestingly, our previous work^22^ pointed to an additional utility of the fast sequential process, which reduces the variability of similar movements, plausibly by reallocating coding resources according to an efficient-coding principle^41,56^. That benefit, however, requires sequential structure and vanishes when targets follow a random sequence. The present results offer a complementary account, showing that the repulsive effect is useful in its own right, reducing the attractive bias even when the target sequence is entirely unpredictable. Notably, the efficient-coding model and the bias-cancellation account predict the same temporal modulation, that autocorrelation should strengthen the sequential effect, because when recent history is more informative, reallocating coding resources yields a greater benefit.

Together, we showed that the motor system relies on two complementary systems. Prior adaptation improves efficiency by exploiting the regularities of a stable context, while the repulsive bias preserves sensitivity to change, safeguarding accuracy when the local context shifts. More broadly, these findings recast movement preparation as an active, predictive process in which the motor system continually reads the marginal and sequential structure of its environment on different timescales and configures its preparatory state to anticipate what is likely to come and quickly correct for recent change. By combining a slowly learned model of the world’s stable structure with a fast correction for its recent changes, the motor system reshapes the speed–accuracy tradeoff rather than being bound by it, preparing movements that are at once fast and accurate.

## Materials and Methods

### Participants

390 young adults (186 female, age: 26.9 ± 4.8 y) were recruited using Prolific.io. Participants performed the experiment on their personal computers through a web-based platform for motor learning experiments. Based on a prescreening survey employed by Prolific, all participants were right-handed and had normal or corrected-to-normal vision. These participants were paid $12/h. All experimental protocols were approved by the Institutional Review Board at the University of California, Berkeley (Approval number: 2016-02-8439). Informed consent was obtained from all participants.

### Task Designs

All experiments were performed using our web-based experimental platform^22,57^. The code was written in JavaScript and presented via Google Chrome, designed to run on any laptop computer. Visual stimuli were presented on the laptop monitor and movements were produced on the trackpad. Data were collected and stored using Google Firebase. Participants were automatically assigned to conditions by the online recruitment platform, Prolific.com. We aimed to recruit around 30 participants for each condition, a standard group size applied in online motor studies^22,52,57^.

#### Experiment 1

Exp 1 (n = 36) was designed to examine whether central tendency effect and repulsive sequential bias can co-exist. To start each trial, participants moved the cursor to a white circle (radius: 1% of the screen height) positioned at the center of the screen. After 500 ms, a red target circle (radius: 1% of the screen height) appeared at a radial distance of 40% of the screen size. At the beginning of the experiment, a center target was randomly selected from 1° to 360° (randomized across participants). On each trial, target locations were randomly generated within a range of −90° to 90° relative to the center target, with a minimum step size of 1°. The red target turned blue after 700 ms, and participants were only allowed to move after the target turned blue. This manipulation forced participants to take time preparing the movement and suppressed target-independent habitual responses. Participants were instructed to produce a rapid shooting movement through the target and to withhold the movement until the target turned blue. Once the movement amplitude reached the target distance, the target disappeared, and no visual feedback was provided. If movement time exceeded 300 ms, the message “Too Slow” was presented. If movement initiation occurred before the target turned blue, the message “Don’t move before target turns blue” was presented. Any warning message disappeared after 1 s. At the end of each trial, the cursor position was reset to a random location within a circle centered at the start position, with a radius equal to 4% of the target distance. Participants then moved the cursor back to the start position to initiate the next trial. The trial was randomized with a cycle of 180 trials, and each participant completed 4 cycles (720 trials).

#### Experiment 2

Exp 2 examined how the marginal statistics of the target distribution reshape the SAT. To probe the SAT function while allowing participants to express their natural response preferences, we applied two response-contingency conditions.

In the delayed-response condition, the procedure was the same as in Exp 1, but we applied two additional target ranges. In the medium-range (n = 31) condition, target locations were randomly generated within a range of −60° to 60° relative to the center target. In the narrow-range (n = 36) condition, target locations were randomly generated within a range of −30° to 30° relative to the center target. The trial was randomized with a cycle of 120 trials for medium-range and 60 trials for narrow-range, and each participant completed 720 trials in total.

In the free-response condition, we removed the enforced delay so that participants could initiate movements at their own pace, where participants tend to hasten their responses at the cost of larger errors. The experimental design was largely identical to that of the delayed-response condition, but with one key difference: after holding at the start position for 500 ms, the blue target appeared, and participants were allowed to initiate their movement immediately upon target onset. We replicated all three prior conditions (wide-range: n = 32; medium-range: n = 29; narrow-range: n = 32).

#### Experiment S1

Exp S1 (n = 64) served as a control experiment for the free-response condition of Exp 2 to quantify the movement preparation time effect that occurred at target detection rather than movement preparation. We applied a simple reaction task. Each trial started with the start circle being presented at the center of the screen. After 500–2500 ms (following an exponential distribution), a blue target appeared at a radial distance of 40% of the screen size. Other visual presentations exactly matched those in the free-response condition. Also, the target sequence was generated in the same way as described in Exp 1 and 2. Participants were instructed to press the space key as soon as they saw the target. After the key press, the target disappeared. If the reaction time was longer than 1 s, we presented the message “Too Slow” for 2 s. After another 1 s, the start circle appeared again, indicating the next trial. We replicated only the medium-range (n = 32) and narrow-range (n = 32) conditions.

#### Experiment 3

Exp 3 examined how the temporal autocorrelation of the target influences the sequential effect. The event sequence within a trial was identical to Exp 1. Participants were randomly assigned to one of three autocorrelation conditions, defined by the average probability that the target changed position from one trial to the next. In the P(switch) = 1 condition (n = 36), a new target was sampled uniformly from −180° to 180° on every trial. In the P(switch) = 1/2 (n = 45) and P(switch) = 1/3 (n = 49) conditions, a target was sampled the same way and then repeated for 1–3 or 1–5 trials, respectively, with the run length drawn uniformly; at the end of each such miniblock a new target was sampled and the process repeated. Participants completed 1080 trials in the P(switch) = 1 condition and 1480 trials in the P(switch) = 1/2 and P(switch) = 1/3 conditions.

### Data Analysis

We computed hand-angle error as the angular difference between the hand position at the target distance and the target position. Trials were excluded if movement duration exceeded 1000 ms or absolute error exceeded 90° (6% of trials). To characterize the systematic bias at the individual level, we fit reaching error as a function of relative target position using a polynomial with a maximum order of 10. The slope of this function and the residual variance served as measures of systematic bias and motor variability, respectively. To isolate the sequential effect, we subtracted the fitted systematic bias from the trial-by-trial reaching error to obtain a motor deviation, and plotted this deviation against the difference in target position between trials *n*−1 and *n* (ΔTarget); the analysis extends to earlier lags (trials *n*−2, *n*−3, …, *n*−10). We summarized the effect with a sequential-effect (SE) index, i.e., the peak-to-peak amplitude of the S-shaped sequential-effect function, computed from the deviation averaged in 20°-wide ΔTarget bins, taken as the mean deviation at the −40° and −20° bins minus the mean deviation at the +20° and +40° bins; for the narrow-range condition, whose targets span a smaller range, the single −20° and +20° bins were used. A positive index indicates repulsion. For Exp 1 and 2, all trials contributed to this analysis. In Exp 3, target directions were locally correlated within miniblocks, which could introduce spurious trial-to-trial dependencies; we therefore computed ΔTarget only for transitions in which trial *n*−*k* was the final trial of a miniblock, a procedure that simulations confirmed recovers the true sequential effect while minimizing within-block contamination (Fig S5). To quantify the within-condition U-shaped modulation of RT (Fig 3d) and MT (Fig S2), we fitted a second-order polynomial to the RT and MT function of target position and tested the quadratic coefficient across participants against zero with a one-sample t-test, where a positive coefficient indicates a U-shape.

Group-level differences in the bias slope, motor variability, reaction time, and SE index were assessed with analysis of variance (ANOVA); homogeneity of variance was checked with Levene’s test, and for groups with unequal variances (Exp 3) we used Welch’s ANOVA with Games–Howell post-hoc comparisons. Effect sizes are reported as partial eta-squared (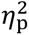). Significance was set at two-tailed p < 0.05.

### Spatial Drift Diffusion Model (sDDM) for prior adaptation

We developed a computational framework to provide a unified account of the observed central tendency bias in reaching behavior and its modulation by reaction time (RT). The model integrates a population-coded stochastic DDM with an experience-dependent prior adaptation mechanism.

#### Evidence Accumulation

The model consists of N = 1000 population units, each tuned to a preferred reaching direction u spanning a circular motor space [0, 360°]. On each trial, the population activity field D starts at an initial level representing the current prior and evolves dynamically through a stochastic evidence accumulation process. The change in activation of the unit tuned to direction u at time t is given by:

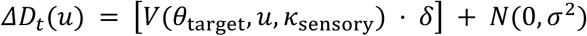

where V(·) denotes the von Mises probability density function, *θ*_target_ is the target direction on the current trial, *κ*_sensory_ controls the precision of sensory encoding, δ is the drift rate governing the speed of evidence accumulation, and N(0, σ²) represents additive Gaussian white noise with variance σ².

A response is initiated when the activation of any unit reaches a decision boundary *B_t_*. Reaction time (RT) is defined as the time step at which this threshold is crossed. The final reaching direction, *θ*_hand_, is determined by the preferred direction of the unit with the maximum activation at decision time. Although the model operates over a circular variable, it is distinct from standard circular DDM formulations in that decision formation emerges from population-level competition rather than a single bivariate diffusion process.

#### Decision Boundary

To account for task-dependent temporal constraints, we implemented distinct decision boundary dynamics across experimental conditions. In the Free-Response Task, to model urgency and the tradeoff between speed and accuracy, we employed a linearly collapsing decision boundary:

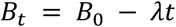

where *B*_0_ is the initial threshold and λ is the boundary collapse rate. In the Delayed-Response Task, rapid responding conferred no benefit. Accordingly, we assumed a fixed decision boundary:

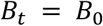

#### Prior Adaptation

Prior adaptation was implemented as a gradual modification of the initial activation field. We assumed that directions encountered more frequently in the environment result in a higher initial state, biasing subsequent decisions. To characterize behavior after learning, we computed the steady-state (balanced) activation field. The expected drive to the initial-state field is expressed as:

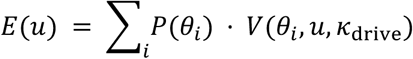

where *κ*_drive_ governs the spatial generalization of prior updating. P(*θ_i_*) is the probability of a target at *θ_i_* given the prior. To simulate Exp 1 and 2, we applied the identical priors as have been implemented empirically. To simulate the experiments with a single repeated target, we applied a single target at θ=0 with P(0)=1.

The steady-state initial state *V*_stable_(u) is derived using a capped delta-rule equilibrium:

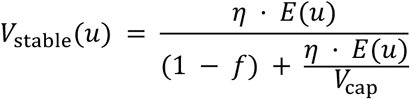

where η is the learning rate, f is a forgetting factor controlling decay toward baseline, and *V*_cap_ imposes a saturation limit on the initial activation to prevent unbounded growth. This steady-state field initializes the accumulation process on subsequent trials and biases both choice and its dependence on RT.

#### Model fitting

We fitted the model to the group-averaged central-tendency function in each condition, because individual data are strongly contaminated by target-specific bias^58^. This is a task-irrelevant source of error that cannot be separated from the central-tendency bias at the individual level but averages out across participants. We first fitted the delayed-response condition, in which the decision boundary is fixed, using the three prior conditions (wide, medium, and narrow) simultaneously. The drift rate (*δ* = 5 × 10^−5^) and the forgetting factor (*f* = 0.1) were held fixed, and six parameters were fitted: the learning rate (*η* = 4.4 × 10^−3^), the concentrations of the two von Mises generalization functions governing evidence accumulation (*κ*_sensory_ = 3.33) and prior adaptation (*κ*_drive_ = 1.14), the decision bound (*B*_0_ = 1.5 × 10^−2^), the saturation cap on the initial-state field (*V*_cap_ = 7.28 × 10^−2^), and the noise (*σ* = 7.25 × 10^−5^). Holding these parameters fixed, we then fitted a single additional parameter, the boundary-collapse rate (*λ* = 8.9 × 10^−5^), to the free-response conditions. Finally, to convert simulated decision times into reaction times, we fitted two further parameters: the duration of a simulation time step (5.0 ms) and a non-decision time (66.3 ms).

### Modeling Sequential Effect

To examine how trial-by-trial sequential dependencies influence overall reaching accuracy in the presence of a central tendency bias, we simulated a generative model in which each response reflects the combined influence of a steady-state central tendency component and a history-dependent sequential bias. On each trial, the produced reaching direction was defined as:

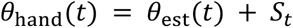

where *θ*_est_(t) captures the central tendency bias measured empirically. Specifically, the central tendency estimate was modeled as a weighted combination of the current target direction *θ_t_* and the participant’s prior mean 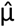:

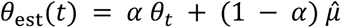

The sequential modulation term *S_t_* was driven by recent target-to-target changes. We defined the target transition as:

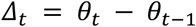

To capture the well-established nonlinear form of sequential effects observed in reaching and perceptual decision tasks^22,59^, *Δ_t_* was transformed through a derivative-of-Gaussian (DoG) sensitivity kernel:

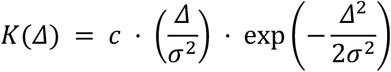

where c determines the overall amplitude of the sequential bias and σ controls its spatial tuning. This kernel has been widely used in previous work to model attractive and repulsive sequential dependencies that peak at intermediate transition magnitudes.

The complete sequential term summed the kernel applied to the one-back transition and a down-weighted (0.1×) contribution from the two-back transition, *S_t_* = *K*(*θ_t_* − *θ_t_*_–1_) + 0.1 · *K*(*θ_t_*_–1_ − *θ_t_*_–2_), with the spatial tuning fixed at σ = 35°. The central-tendency weight α was not fit freely but set so that the model reproduced the empirically measured attractive bias in each condition (α = 1 − bias slope).

To evaluate how sequential effects influence global task performance, we systematically varied the sequential gain parameter c of the DoG kernel over a range of values (c = 1–20). For each value, we generated simulated reaching responses using the same target sequences applied to each participant within a given prior condition (or switch-probability condition). For Exp 3, we assumed no central tendency component ( *α* = 1), because targets were uniformly sampled from the full circle, eliminating a stable directional prior. Performance was quantified using the exact same method as the empirical data.

#### Reanalysis of use-dependent learning data

We reanalyzed a previously published UDL dataset^23^ to test whether the sDDM accounts for the coupling between RT and reaching error in this environment. Their Exp 1 and Exp 2 correspond to our free-response and delayed-response conditions, respectively. Participants reached toward a repeated target on 90% of trials; on the remaining 10%, a probe target displaced from the repeated target was presented. For illustration, we plotted reaching error for the 90° probe target as a function of RT, pooled across participants (Fig 4a). The sequential effect in the UDL design (Fig 6b) was measured on repeated-target trials that immediately followed a probe trial. Percent compensation (Fig 6c) quantifies how much of the attractive central-tendency bias is canceled by the repulsive sequential effect. For each target position we computed the predicted attractive bias and the sequential effect expected from the preceding target. Because the observed bias is already reduced by the sequential effect, the underlying attractive bias was recovered as the sum of the two, and compensation was defined as the fraction of that underlying bias offset by the sequential effect: 100% × |Σ*SE*| / (|Σ*bias*| + |Σ*SE*|). We computed compensation for the delayed-response, wide-prior condition of Exp 2, which served as the random-target condition, and for the delayed-response condition of the UDL dataset.

## Acknowledgments

TW is funded by C.V. Starr Fellowship. DW is funded by NIH R01CA236793.

## Author contributions

T.W. and D.W. conceived and designed the study. T.W. implemented the experiments, collected and analyzed the data, and developed the computational model. T.W. and D.W. interpreted the results and wrote the manuscript.

## Supplementary Information

**Figure S1.**
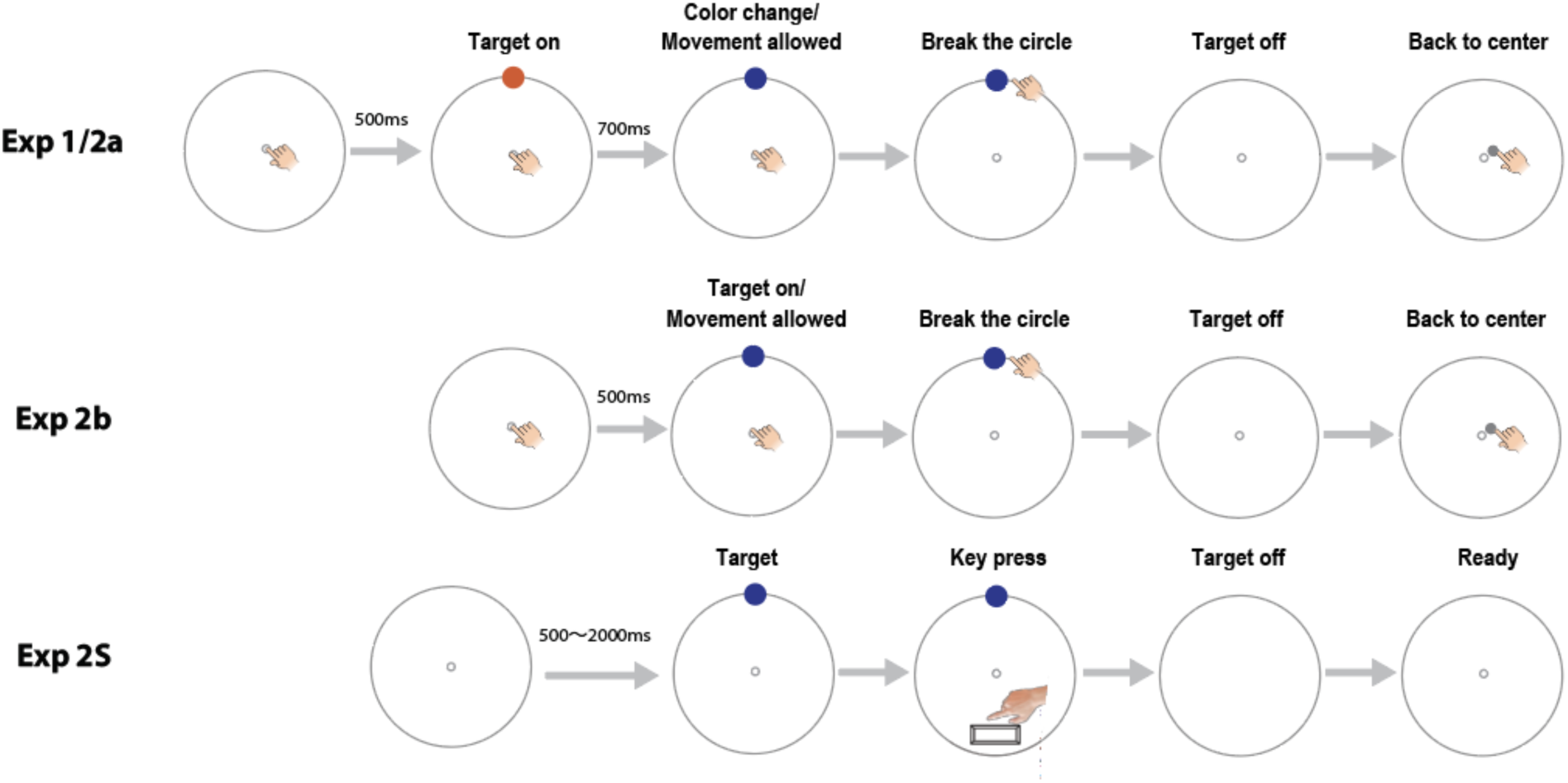
Trial event sequence in Exp 1–2. In the delayed-response task, the target turned blue (the go cue) 700 ms after onset, enforcing full preparation, whereas in the free-response task, the target appeared blue at onset, allowing immediate, self-paced initiation. The visual detection control (Exp S1) replicated the trial sequence of the free-response task, but participants pressed a key as soon as they detected the target rather than making a reach. To ensure responses reflected visual detection rather than a timed key press, target onset time was jittered in this control.

**Figure S2.**
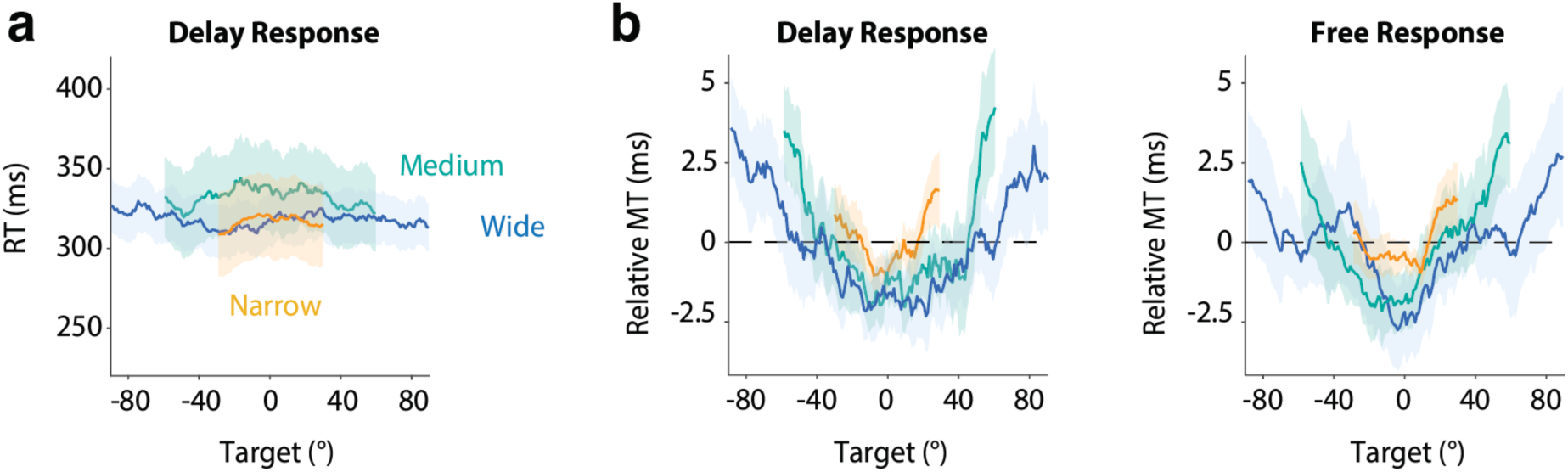
Target distribution modulates movement kinematics. (a) Given the delay period preceding the go cue (see Fig S1), reaction times remain relatively consistent across different target locations and prior conditions in the delayed-response condition. (b) In contrast, movement times exhibit a U-shaped profile with prior-dependent modulation in both response conditions, mirroring the RT modulation in the free-response condition.

**Figure S3.**
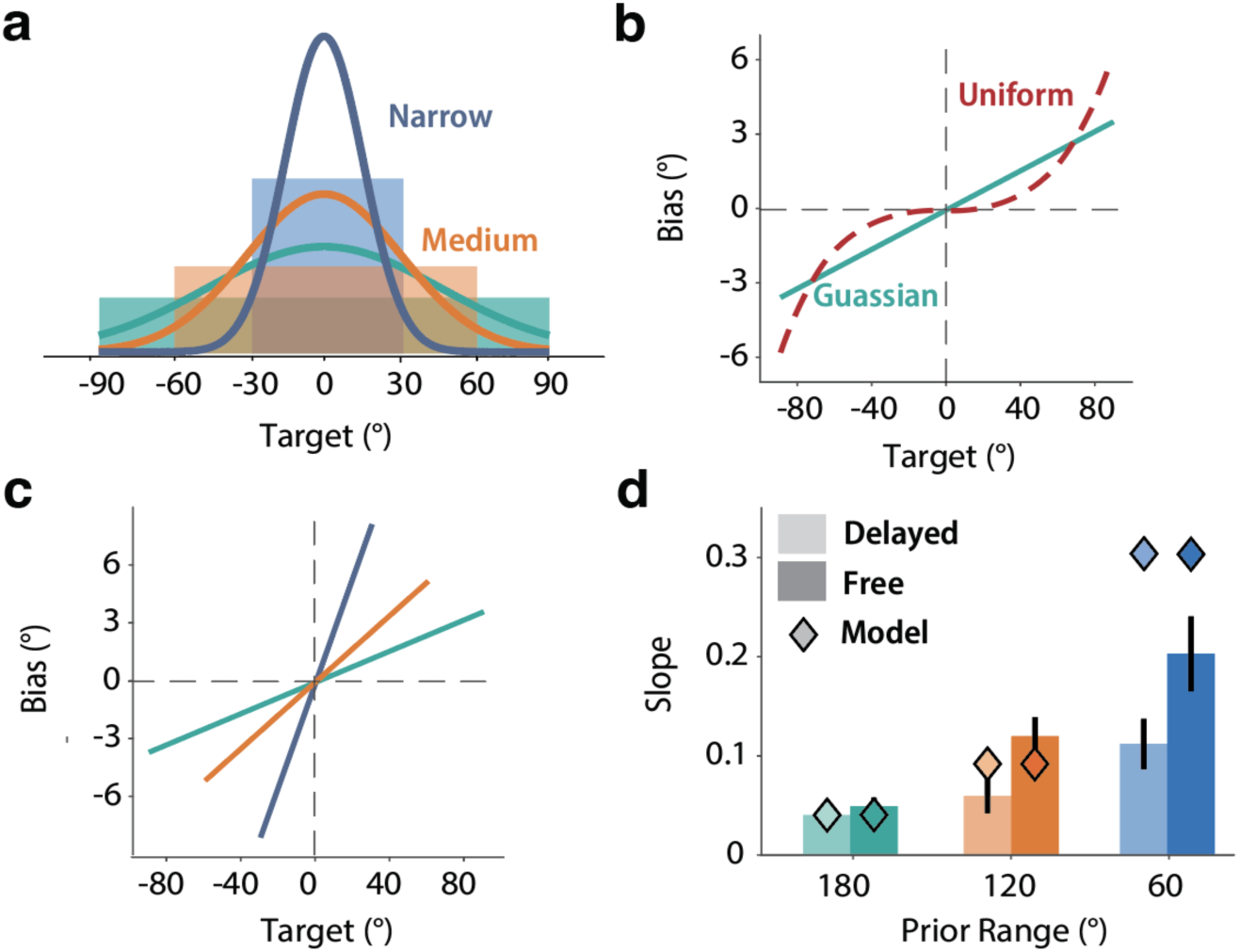
The central-tendency bias does not follow a Bayesian inference model. (a) Prior distributions used in Exp 2, together with Gaussian approximations matched in mean and standard deviation. (b) Central tendency in the wide-prior condition predicted by a Bayesian model assuming the internal prior is either uniform (the ground truth) or a Gaussian approximation of the ground truth. A uniform prior produces an S-shaped bias function, whereas a Gaussian prior produces a linear bias function, closer to what is observed. (c) Central-tendency functions across the three prior distributions predicted by a Bayesian model with a Gaussian-approximate prior. The model predicts the maximal bias increases dramatically as the prior range decreases. (d) Predicted slope of the central tendency function from the Bayesian model across prior conditions and response contingency, plotted on top of the data. The Bayesian model strongly overpredicts how much the central-tendency bias increases as the prior narrows, which is particularly salient in the narrow range condition. Moreover, the model has no mechanism to produce the difference between the two response types.

**Figure S4.**
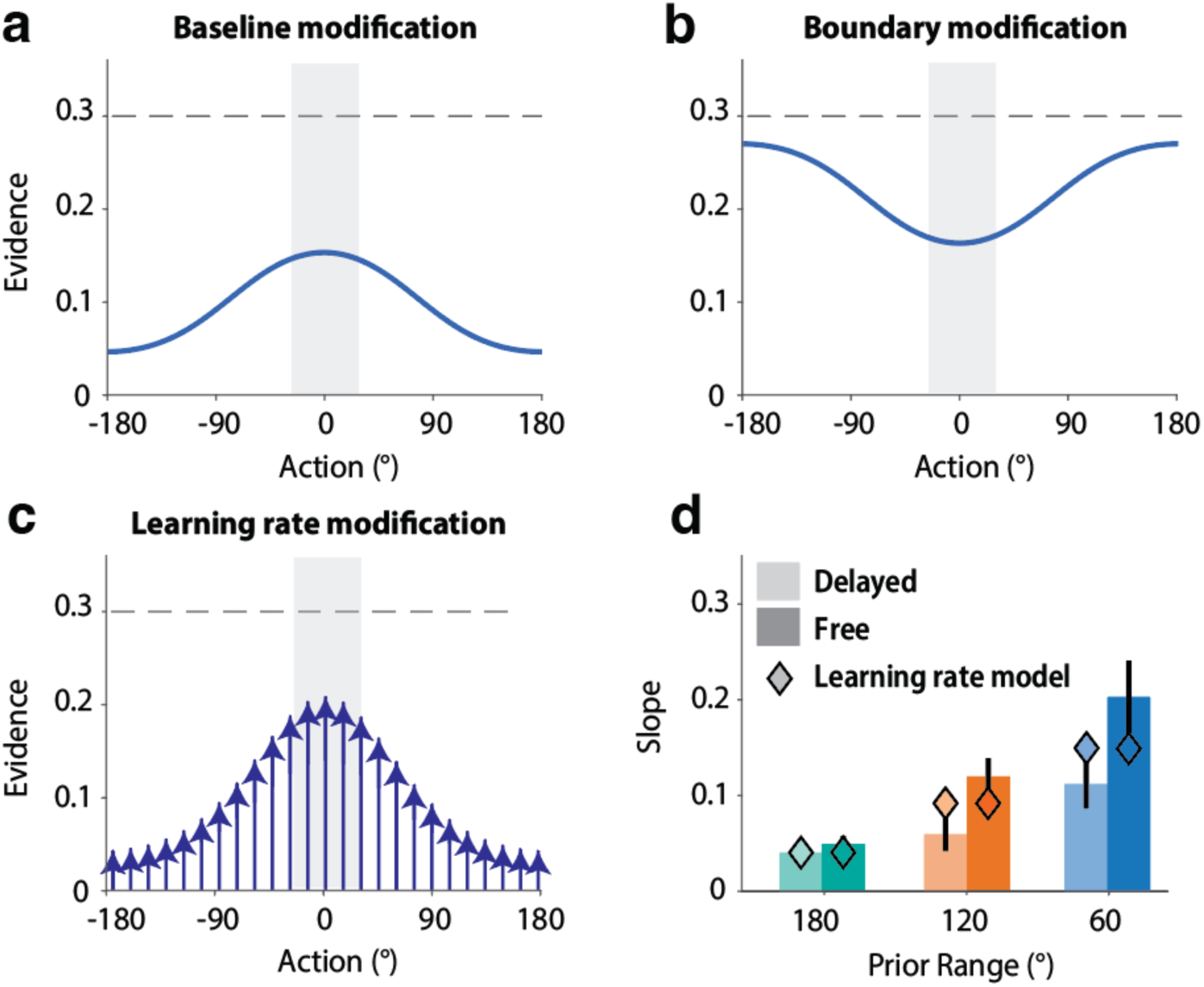
Three candidate mechanisms for prior-driven bias in the spatial drift–diffusion model. (a) Initial-state modification: the prior raises the initial preparation state for actions near the prior mean, so less additional evidence is required to select them. (b) Boundary modification: the prior lowers the decision boundary near the prior mean (blue) relative to the unmodified boundary (dashed). This model is computationally identical to (a), serving as an alternative mechanism. (c) Learning-rate modification: the prior increases the rate at which evidence accumulates for actions near the prior mean (arrows), leaving the initial state and boundary unchanged. (d) Central-tendency slope for the delayed-response and free-response conditions across the three prior ranges, with the learning-rate model’s prediction. The learning-rate model predicts that the central tendency strengthens as the prior range narrows, but it cannot predict the stronger central tendency in the free-response condition than in the delayed-response condition.

**Figure S5.**
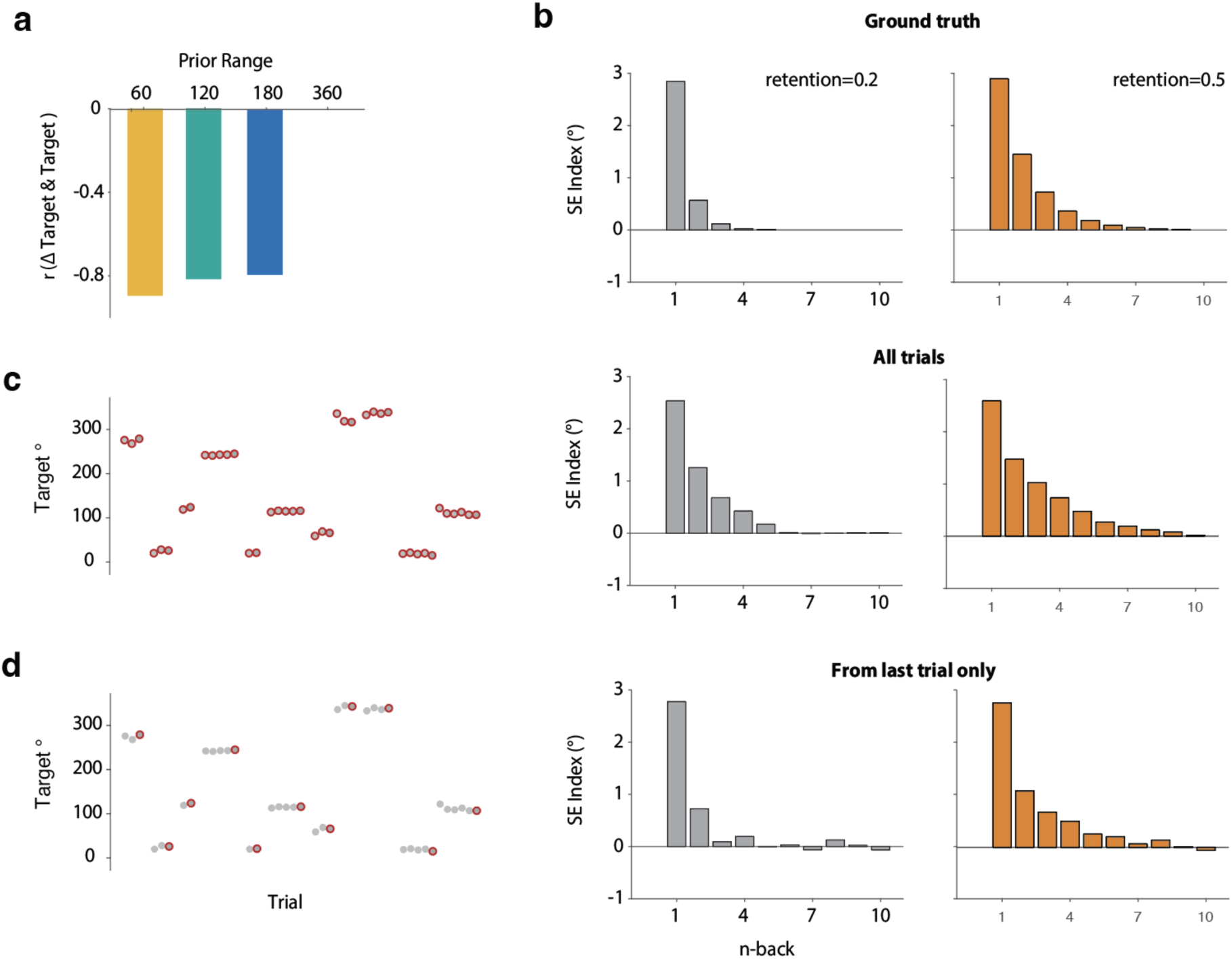
Model simulations reveal appropriate analytical methods for measuring sequential effects. (a) ΔTarget is highly correlated with the current Target position when the distribution is non-circular (i.e., less than a 360° range). This correlation influences the partitioning of the central tendency and the sequential effect (see Fig 5f). (b) Ground truth of the sequential effects applied in the simulations. We tested two different retention rates, where the sequential effect decays at different speeds. (c) If the sequential effect is analyzed using all trials from Exp 3, the recovered effect suffers from an artifact that overestimates the retention rate. (d) If the analysis is restricted to the sequential effect originating from the last trial of each miniblock, the recovered effect precisely reflects the ground truth. Based on these simulation results, we adopted this procedure for analyzing the empirical data in Exp 3.

